# Direct control of somatic stem cell proliferation by the *Drosophila* testis stem cell niche

**DOI:** 10.1101/110619

**Authors:** Eugene A. Albert, Olga A. Puretskaia, Nadezhda V. Terekhanova, Christian Bökel

**Affiliations:** Centre for Regenerative Therapies Dresden, Technical University Dresden, Fetscherstr. 105, 01307 Dresden, Germany; Sector for Molecular Evolution, Institute for Information Transmission Problems of the RAS (Kharkevich Institute), Moscow 127994, Russia; N. K. Koltsov Institute of Developmental Biology of the RAS, Moscow 119334, Russia; Laboratory of Molecular Genetics, Russian Federal Research Institute of Fisheries and Oceanography, Moscow 107140, Russia

**Keywords:** DamID, Kibra, Salvador, stem cell niche, Zfh1

## Abstract

Niches have traditionally been characterized as signalling microenvironments that allow stem cells to maintain their fate. This definition implicitly assumes that the various niche signals are integrated towards a binary fate decision between stemness and differentiation. However, observations in multiple systems have demonstrated that stem cell properties such as proliferation and self renewal can be uncoupled at the level of niche signalling input, which is incompatible with this simplified view. We have studied the role of the transcriptional regulator Zfh1, a shared target of the Hedgehog and Jak/Stat niche signalling pathways, in the somatic stem cells of the *Drosophila* testis. We found that Zfh1 binds and downregulates *salvador* and *kibra*, two tumour suppressor genes of the Hippo/Wts/Yki pathway, thereby restricting Yki activation and proliferation to the Zfh1 positive stem cells. These observations provide an unbroken link from niche signal input to an individual aspect of stem cell behaviour that does not, at any step, involve a fate decision. We discuss the relevance of our observations and other reports in the literature for an overall concept of stemness and niche function.

**Summary statement:** We demonstrate that the fly testis niche controls stem cell proliferation by repressing Hippo pathway genes independent of a binary cell fate decision between stemness and proliferation.

## Introduction

Stem cell proliferation and maintenance are typically regulated by niches, signalling microenvironments within tissues that allow resident stem cells to fulfill their functions (Scadden, 2014). Since the first niches were characterized at the molecular level in the *Drosophila* ovary (Xie and Spradling, 2000) and testis (Kiger et al., 2001; Tulina and Matunis, 2001), niche based regulation of stem cells has been observed in a wide variety of systems, and has largely been discussed using a conceptual framework that considers stemness and differentiation as alternative outcomes of a cell fate decision. Implicitly, this definition suggests that a stem cell makes a choice between retaining stemness or commiting to differentiation by integrating the incoming niche signals. Niches were accordingly seen as a means by which the organism can influence this binary decision (Fuller and Spradling, 2007). However, this simplified model is at odds with recent observations in various model system, including again the *Drosophila* gonadal stem cell niches. In the fly testis, stem cell proliferation and self-renewal were shown to be genetically separable at the level of Hh niche signalling input (Amoyel et al., 2013; Michel et al., 2012). Similarly, experimentally altering MAPK activity can uncouple stem cell self-renewal from the regulation of stem cell competitiveness (Amoyel et al., 2016; Singh et al., 2016). These and other observations raise fundamental questions about the nature of stemness and the function of niches.

The *Drosophila* testis offers an excellent model system to address such issues, as niche signalling activity can be genetically manipulated and the outcome monitored with single cell resolution. At the tip of the testis, two types of stem cells, the germline stem cells (GSCs) and the somatic cyst stem cells (CySCs), surround a group of postmitotic, somatic niche cells termed hub that provides multiple niche signals regulating both stem cell pools. In the somatic lineage, the CySCs are the only dividing cells, giving rise to differentiating cyst cells (CyCs) that exit the niche with the developing germline clusters (Losick et al., 2011). CySC maintenance and proliferation requires niche signals by Unpaired (Upd) (Leatherman and Dinardo, 2008) and Hedgehog (Hh) (Amoyel et al., 2013; Michel et al., 2012). Both ligands are produced by the hub and activate the Jak/Stat and Smoothened signalling cascades, respectively, within the CySCs. Even though the two pathways each have multiple, specific targets, they converge on the large, zinc finger and homeodomain containing transcription factor Zfh1 (Fortini et al., 1991). Zfh1 marks the somatic CySC pool within the testis and is absent from differentiated CyCs (Leatherman and Dinardo, 2008) (Figure 1A). Similar to clonal inactivation of either Hh or Upd signalling, removal of their shared target *zfh1* leads to the loss of the mutant CySCs from the stem cell pool by differentiation (Amoyel et al., 2013; Leatherman and Dinardo, 2008; Michel et al., 2012). Zfh1 is thus required for CySC stemness.

**Figure 1.**
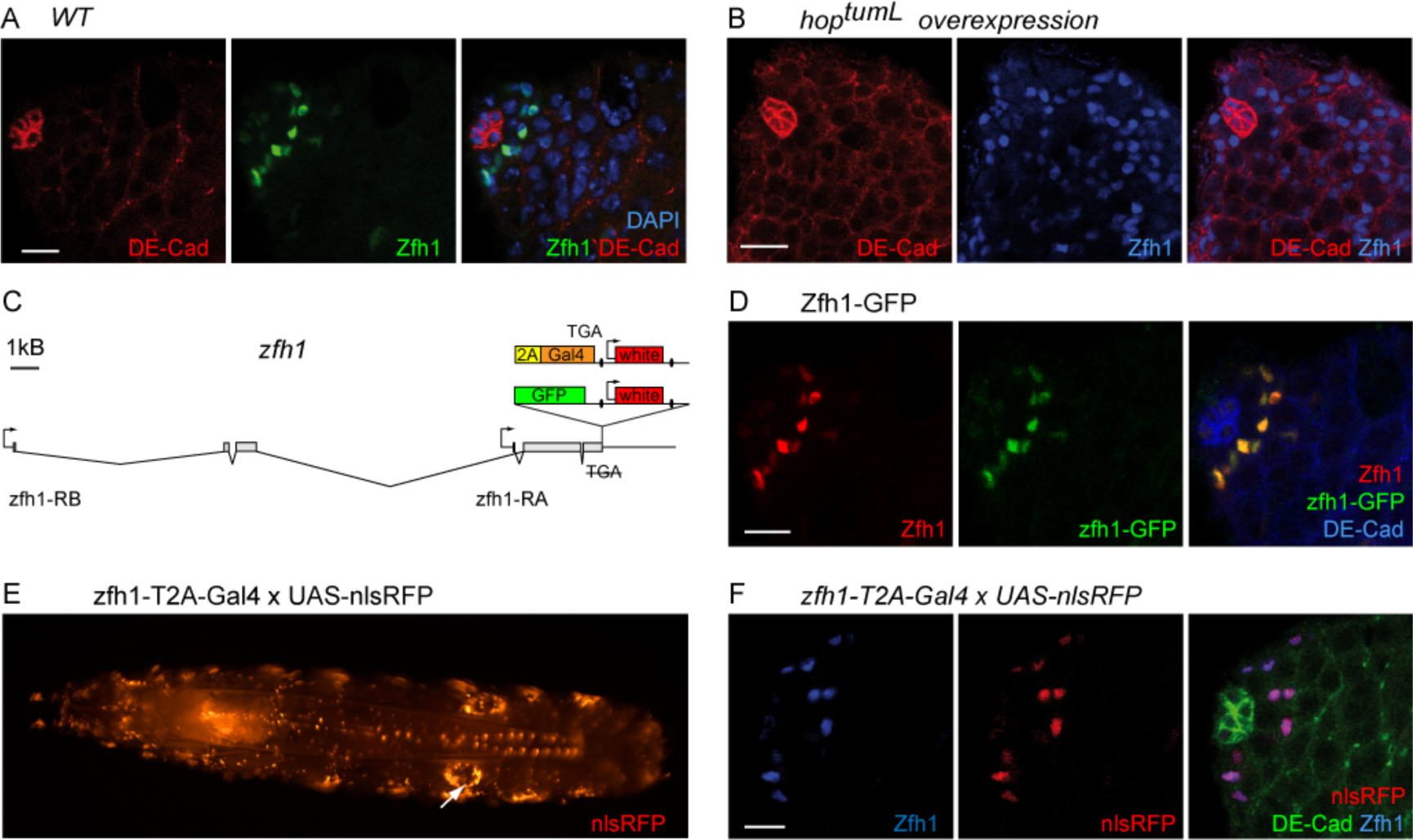
Endogenous Zfh1 expression and Crispr/Cas9 generated reagents. **A** CySCs are recognized by Zfh1 positive nuclei (green) one tier removed from the hub (DE-Cad, red). Nuclei marked by DAPI (blue). **B** Zfh1 positive cells (blue) expand following a 2d pulse of the constitutively active Jak kinase Hop^tuml^. Hub and CyC outlines marked by DE-Cad (red).**C** Schematic representation of the Zfh1 locus and C-terminal GFP and T2A-Gal4 fusions. Ellipses indicate loxP sites. **D** Zfh1-GFP (green) colocalizes with Zfh1 immunostaining (red) in the vicinity of the hub (DE-Cad, blue). **E,F** nlsRFP expression under control of zfh1-T2A-Gal4. **E** In the larva, RFP is expressed in multiple tissues including the testis (arrow). **F** in the adult testisnlsRFP (red) coincides with Zfh1 protein (blue) labelling the CySCs abutting the hub (DE-Cad, green). Scale bars 10*μ*m.

It has been proposed that Zfh1 expression is also sufficient for CySC proliferation and self-renewal (Leatherman and Dinardo, 2008). However, the evidence for this is less clear: Upd overexpression or Jak/Stat pathway overactivation causes the proliferation of Zfh1 positive, CySC-like cells (Leatherman and Dinardo, 2008) (Figure 1B) that retain some stem cell characteristics such as proliferation and non-differentiation (Leatherman and Dinardo, 2008) but also acquire certain niche properties including the expression of BMP ligands (Leatherman and Dinardo, 2010). This renders them able to recruit and maintain GSCs, a niche activity endogenously restricted to the hub (Inaba et al., 2015; Kawase et al., 2004; Michel et al., 2011), resulting in stably proliferating, mixed lineage tumours. Prolonged, Gal4 mediated overexpression of Zfh1 indeed causes a similar phenotype. However, while increased CySC proliferation in response to Zfh1 overexpression is rapidly detectable, tumour formation resembling the Jak/Stat pathway overactivation phenotype only becomes observable after weeks of overexpression (Leatherman and Dinardo, 2008), suggesting that additional regulatory events or signalling inputs must be involved.

Furthermore, overactivation of the Hh pathway, e.g. by inactivation of *patched* (*ptc*), also upregulates Zfh1 expression and causes accelerated proliferation of the mutant clones (Amoyel et al., 2013; Michel et al., 2012). However, despite initially expressing high levels of Zfh1, the affected cells retain the ability and propensity to differentiate (Amoyel et al., 2013; Michel et al., 2012), again suggesting that Zfh1 expression alone is not sufficient for specifying a stable CySC fate.

We therefore hypothesized that the primary role of Zfh1 was to promote CySC proliferation in response to either niche signal, and decided to identify Zfh1 target genes by mapping its binding sites in the CySCs through an *in vivo* DamID approach (Southall et al., 2013).

Here we report that i) Zfh1 binds to the putative control regions of multiple genes encoding upstream components of the Hippo/Warts/Yki signalling cascade, including the tumour suppressors *sav* and *kibra*, that ii) Zfh1 *in vivo* partially suppresses the transcription of both genes, and that iii) activation of Yki, which is necessary and sufficient for stem cell proliferation, is restricted to the Zfh1 positive CySCs. Yki-mediated proliferation of the somatic stem cells in the fly testis can therefore be traced back all the way to the Hh and Upd niche signalling input.

Importantly, none of the steps from the niche ligands to nuclear Yki appears to constitute a cell fate decision establishing stemness. Our observations thus provide a mechanistic explanation for the direct and independent regulation of stem cell proliferation by the niche. We discuss the implications of these and other recent results for the overall concept of stemness, and propose a new, “micromanagement” model of niche function, whereby multiple niche signals directly and independently control distinct subsets of stem cell behaviour.

## Results

### C-terminally tagged Zfh1 fusion proteins are functional

Visualization of expression patterns by GFP reporters and temporally and spatially controlled expression using the Gal4 system are key elements of the *Drosophila* toolbox. However, both were still missing for *zfh1* when we started studying how this gene links niche signals and CySC proliferation. The *zfh1* locus possesses two transcription starts separated by about 17kb. The respective transcripts are spliced to common 3′ exons (Figure 1C), giving rise to two isoforms that are both expressed in the testis (Figure S1). *zfh1-RA* encodes a 747aa protein containing seven C2H2 zinc fingers and a central homeodomain, while the 1045aa protein encoded by *zfh1-RB* contains two additional, N-terminal zinc fingers. We therefore targeted GFP to the common C-terminus of both isoforms (Figure 1C) by Crispr/Cas9 mediated recombination (Gratz et al., 2014; Port et al., 2014). The resulting Zfh1-GFP knockin lines were homozygous viable and fertile. In the testis, Zfh1-GFP marks the nuclei of somatic cells in the first tier surrounding the hub, coinciding with the expression of endogenous Zfh1 in the CySCs (Figure 1D).

To generate a driver line that captures expression of both isoforms we fused Gal4 to the Zfh1 C-terminus via a cotranslationally separating T2A peptide (Szymczak et al., 2004) (Figure 1C). The zfh1-T2A-Gal4 knockin flies were subviable in the homozygous state (ca. 60% of expected adults hatching) and largely male sterile over a deficiency deleting the *zfh1* locus. Viability improved to Mendelian ratios and fertility recovered when the *w*^+^ transgenesis marker was excised by cre/lox recombination. In the larva, zfh1-T2A-Gal4 driven expression of nls-RFP uncovered a complex expression pattern that encompassed multiple tissues with reported Zfh1 presence (Broihier et al., 1998; Lai et al., 1991), including e.g. musculature, heart, nervous system, hemocytes, and gonads (Figure 1E). In the adult testis, RFP expression under control of zfh1-T2A-Gal4 was confined to the endogenously Zfh1 positive CySCs (Figure 1F), demonstrating that the knockin line can direct transgene expression to our cells of interest.

Zfh-1 can therefore in principle tolerate C-terminal fusions without affecting protein function. Since the existing UAS-Zfh1 constructs (Postigo and Dean, 1999) are based on *zfh1-RB*, we opted for C-terminal modification of this isoform for the cDNA based DamID constructs described below.

### Zfh1 binding sites identified by DamID in S2 cells are enriched at active regulatory regions

To identify Zfh1 target sites in the genome we chose a NGS based DamID approach based on the TaDa system (Southall et al., 2013) that uses an mCherry leader ORF termed LT3 to achieve the desired, low expression levels of the Dam and Dam fusion proteins (Figure S2A). We first validated this approach for Zfh1 by mapping its binding to the DNA of cultured S2 cells that also endogenously express Zfh1 (Figure S2B). We generated stably transfected, polyclonal S2 cell lines expressing either LT3-Dam or LT3-Zfh1-Dam under control of the metallothionein promoter. Applying the damid_seq analysis pipeline (Marshall and Brand, 2015) to our samples flagged 1125 peaks (Figure 2A) associated with 1052 genes based on two replica experiments (Spearman correlation between replicates ᵨ = 0.95 (dam) and ᵨ = 0.96 (zfh1-dam) (Figure 2B). Using a resampling / permutation approach (Zhu et al., 2010) we found that the S2 Zfh1 DamID peaks exhibited significant overlap with Zfh1 binding sites previously identified by ChIPseq in Kc167 cells (Negre et al., 2011) (n=134, permutation test, p<0.001) (Figure 2C). Significant overlap was also observed between the genes associated with the Zfh1 peaks in either cell type (n=179, χ^2^-test, p<0.0001) (Figure 2D). In addition, 70% of the Zfh1 DamID peaks we identified in S2 cells coincided with regions flagged as enhancers in the same cell type by STARR-seq (Arnold et al., 2013) (Figure S2C), again significantly more than expected by chance (n=785, permutation test, p<0.001) (Figure 2E).

**Figure 2.**
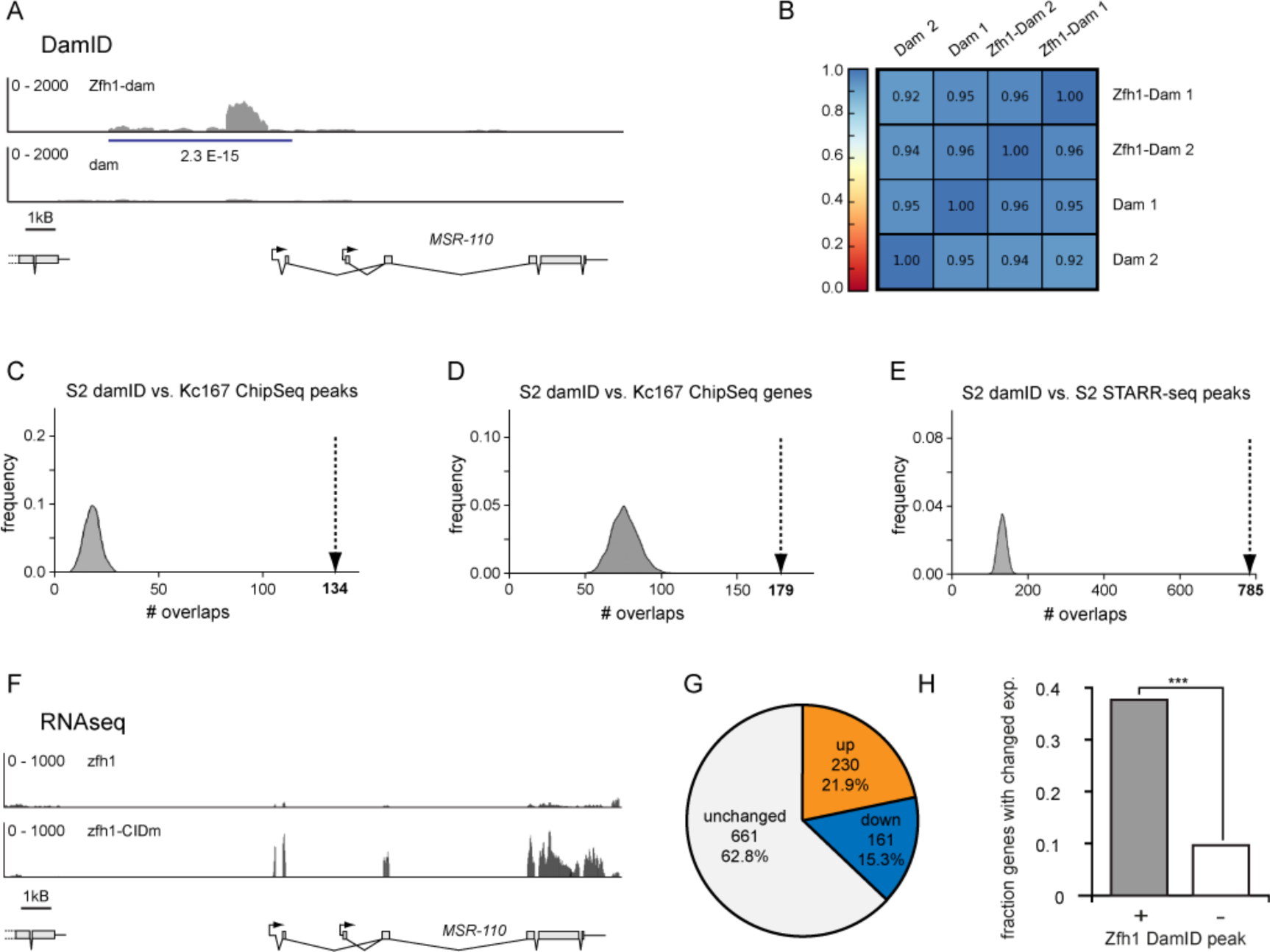
Zfh1 DamID in S2 cells. **A** NGS reads from a Zfh1-Dam sample (top panel) are enriched relative to a Dam only control (bottom panel) near the MSR-110 distal transcription start (peak as called by damid_seq, blue line; FDR indicated). **B** Spearman correlation between replica experiments. **C** Zfh1 DamID peaks in S2 cells colocalize with Zfh1 ChIP peaks in Kc167 cells (p<0.001, permutation test). Distribution of overlaps from 1000 resamplings plotted against frequency; dashed arrow indicates observed value. **D** Same as **C** for the respective associated genes (p<0.001, χ^2^-test). **E** Zfh1 DamID peaks overlap (p<0.001, permutation test) with enhancer regions identified by STARR-seq. **F-H** Transcriptional changes following expression of a Zfh1 construct unable to bind CtBP (zfh1-CIDm). **F** MSR-110 exhibits transcriptional derepression. **G** Response to zfh1-CIDm expression by genes associated with a Zfh1 DamID peak. **H** Genes associated with a Zfh1 DamID peak are significantly more likely to exhibit changed transcription levels (***, p<0.001, χ^2^-test).

Finally, we compared the transcriptomes of S2 cells overexpressing either functional Zfh1 or a Zfh1 version unable to bind the transcriptional corepressor CtBP (Zfh1-CIDm) (Postigo and Dean, 1999). Upon overexpression of Zfh1-CIDm, 230 out of all 1052 genes associated with Zfh1 DamID peaks (21.9%) exhibited the expected signature of transcriptional derepression (Figure 2F,G). A comparable number instead showed decreased transcription levels (161/1052; 15.3%) (Figures 2G and S2D), while the majority of genes associated with a Zfh1 peak experienced no significant change (661/1052 genes; 62.8%) (Figures 2G and S2E). Zfh1 may therefore, like its mammalian homologue ZEB1 (Gheldof et al., 2012), not exclusively act as a CtBP-dependent transcriptional repressor. Importantly, though, the fraction of genes exhibiting changes in transcription in either direction was significantly higher amongst genes associated with at least one Zfh1 peak than for the overall transcriptome (37.5% vs. 9.3%, χ^2^-test: p<0.0001) (Figure 2H). Thus, DamID using a Zfh1-Dam fusion protein uncovered binding sites enriched for functional, Zfh1-dependent regulatory elements.

### Identifying Zfh1 target genes *in vivo*

However, while these results demonstrated that DamID can in principle uncover Zfh1 binding sites, there was no way of predicting *a priori* which of the targets identified in cell culture may be relevant in the testis niche *in vivo*. We therefore decided to use the same fusion protein for mapping Zfh1 binding sites directly in the CySCs of the adult testis, and generated transgenic flies expressing Zfh1-Dam with the LT3 mCherry leader ORF under UAS control (UAS-LT3-zfh1-Dam) (Figure S3A). To suppress transgene expression prior to induction we recombined tub-Gal80^ts^ transgenes both onto the chromosomes carrying the UAS-LT3-dam or UAS-LT3-Zfh1-dam insertions and onto the zfh1-T2A-Gal4 driver line. Following a 24h pulse of transgene expression DNA was then extracted from testes of adult males of the appropriate genotypes.

Applying the same analysis as above uncovered 811 Zfh1 peaks (Figure 3A) present in both replicates (Spearman correlation between replicates ᵨ = 0.97 (dam) and ᵨ = 0.90 (Zfh1-dam)) (Figure 3B). 377 of these sites (46%) were also occupied in S2 cells (Figure 3C), corresponding to 420 of the 968 genes associated with Zfh1 in CySCS (43%) (Figure 3D). 18% of the CySC zfh1 DamID peaks, again significantly more than expected by chance (n=150, permutation test, p<0.001), coincided with enhancers identified by STARR-seq in cultured ovarian somatic cells (OSCs) (Arnold et al., 2013), mesodermal cells developmentally related to the CySCs (Figure 3E,F). Significant overlap was also observed between the CySC Zfh1 DamID peaks and the Kc167 Zfh1 ChIP peaks (n=67, permutation test, p<0.001) and between the associated genes (n=113, χ^2^-test, p<0.001) (Figure S3B,C). Finally, Gene Ontology analysis using Panther GO-Slim (Mi et al., 2016) revealed that a limited number of biological processes (7 of 222 terms) was overrepresented among the putative Zfh1 target genes (Figures 3G and S3D), amongst which the GO term “signal transduction” immediately raised our interest as a potential link to CySC proliferation.

**Figure 3.**
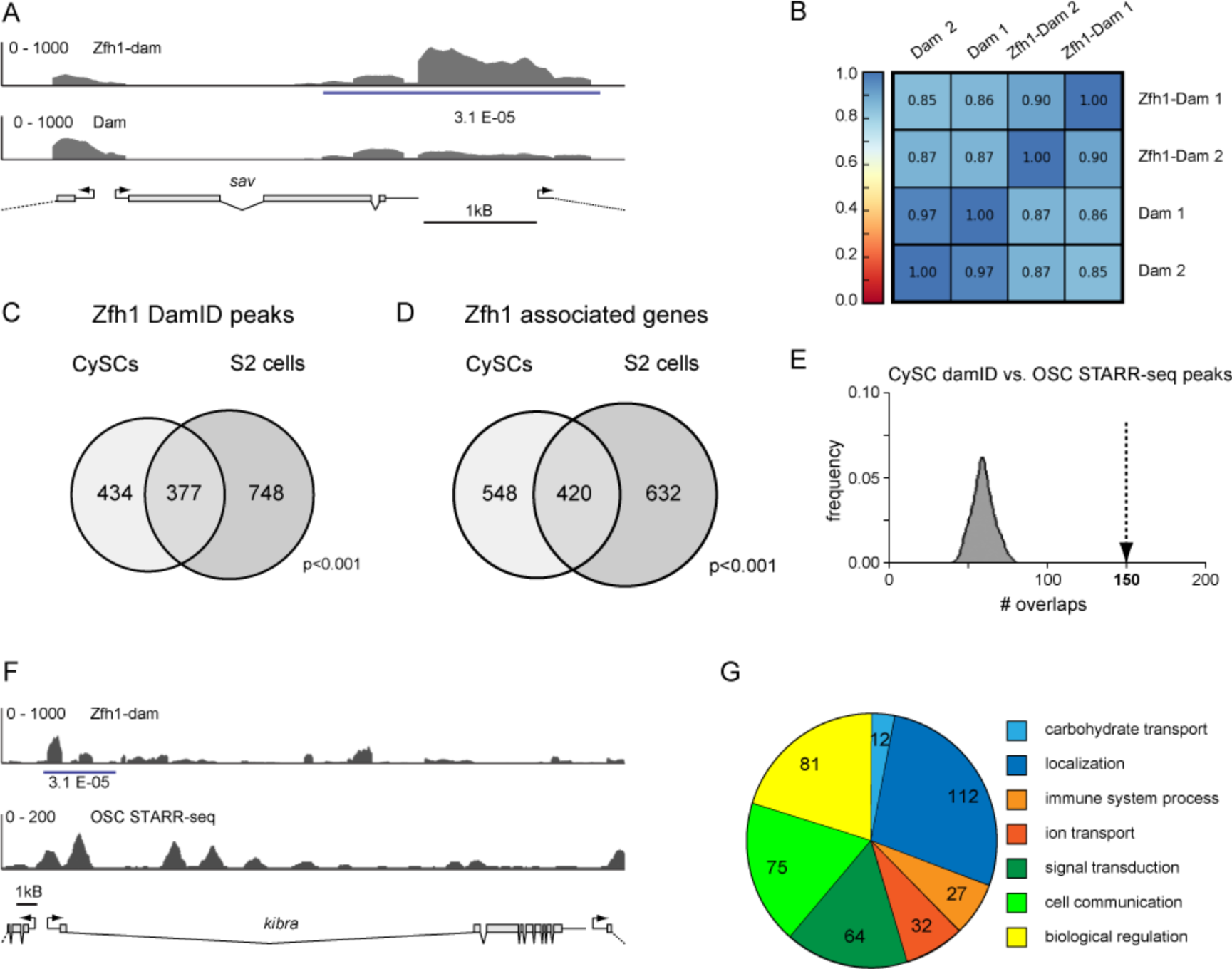
*In vivo* Zfh1 DamID in CySC cells. **A** Zfh1 DamID reads are enriched near the 3’-UTR and intergenic region of the *sav* locus (extent of DamID peak, blue line; FDR indicated). **B** Spearman correlation between replica experiments. **C,D** 46% of CySC DamID peaks **C** and 43% of the associated genes **D** are also occupied in S2 cells, significantly more than expected by chance (C, p<0.001, permutation test; D, p<0.001, χ^2^-test). **E** CySC Zfh1 DamID peaks overlap with ovarian somatic cell enhancer regions (p<0.001, permutation test) **F** Comparison of Zfh1 damID peaks and OSC STARR-seq peaks for the *kibra* locus **G** GO-slim terms enriched among Zfh1 associated genes.

### Zfh1 binds at or near genes encoding members of the Hippo signalling cascade

Since our aim was to understand the link between niche activity and stem cell proliferation we were intrigued by a cluster of hits in or near genes encoding components of the Hippo signalling cascade. This pathway is generally associated with cell growth, proliferation, and survival (Enderle and McNeill, 2013; Irvine and Harvey, 2015; Meng et al., 2016), and had been shown to be necessary and sufficient for controlling CySC proliferation (Amoyel et al., 2014). However, it remained unknown whether or how Hippo pathway activity was regulated by niche signals.

The Hippo signalling cascade consists of a core of two serine/threonine kinases, Hippo (Hpo) and Warts (Wts), that phosphorylates and thus inactivates the effector transcription factor Yki. Activity of the core complex depends on the associated adapter proteins Salvador (Sav) (Tapon et al., 2002) and Mob-as-tumour-suppressor (Mats), and on scaffolding proteins such as Expanded, Kibra (Baumgartner et al., 2010; Genevet et al., 2010; Yu et al., 2010), and Merlin that recruit the complex to the plasma membrane. Inactivation of the upstream components or the kinase complex, either by physiological signals or by mutation, allows Yki to enter the nucleus and promote target gene expression. In CySCs, we detected Zfh1 binding at or near the *kibra*, *sav*, and *mats* genes (Figures 3A and S4). In S2 cells the peak near *mats* was absent. Instead, binding was additionally observed at *ex*, *pez*, *wts*, and *sd* (Figure S4).

The ability of overexpressed Zfh1 to induce somatic cell proliferation in the testis depends on its function as a CtBP-dependent transcriptional repressor (Leatherman and Dinardo, 2008; Postigo and Dean, 1999). Binding of Zfh1 to Hpo pathway tumour suppressors thus immediately suggested a direct and linear connection between niche signalling and stem cell proliferation: Upd and/or Hh niche signals produced by the hub may locally activate Zfh1 expression in adjacent somatic cells, which would in turn repress multiple Hpo pathway components, thereby activating Yki exclusively in the Zfh1 positive stem cells. Consistent with this model, a transcriptional reporter for Yki activity (diap1-GFP4.3), in which a minimal Yki response element derived from the *diap1* gene drives expression of a nuclear GFP (Zhang et al., 2008), is in the adult testis largely restricted to the Zfh1 positive CySCs (89±9% of all Zfh1 positive cells and 84±7% diap-GFP positive cells double positive) (Figure 4A). Moreover, Zfh1 overexpression in the somatic lineage using the traffic jam (Tj)-Gal4 driver expanded the region exhibiting Yki activation (Figure 4B,C), while the associated increase in somatic cell count was sensitive to *yki* copy number (Figure S5A,B).

**Figure 4.**
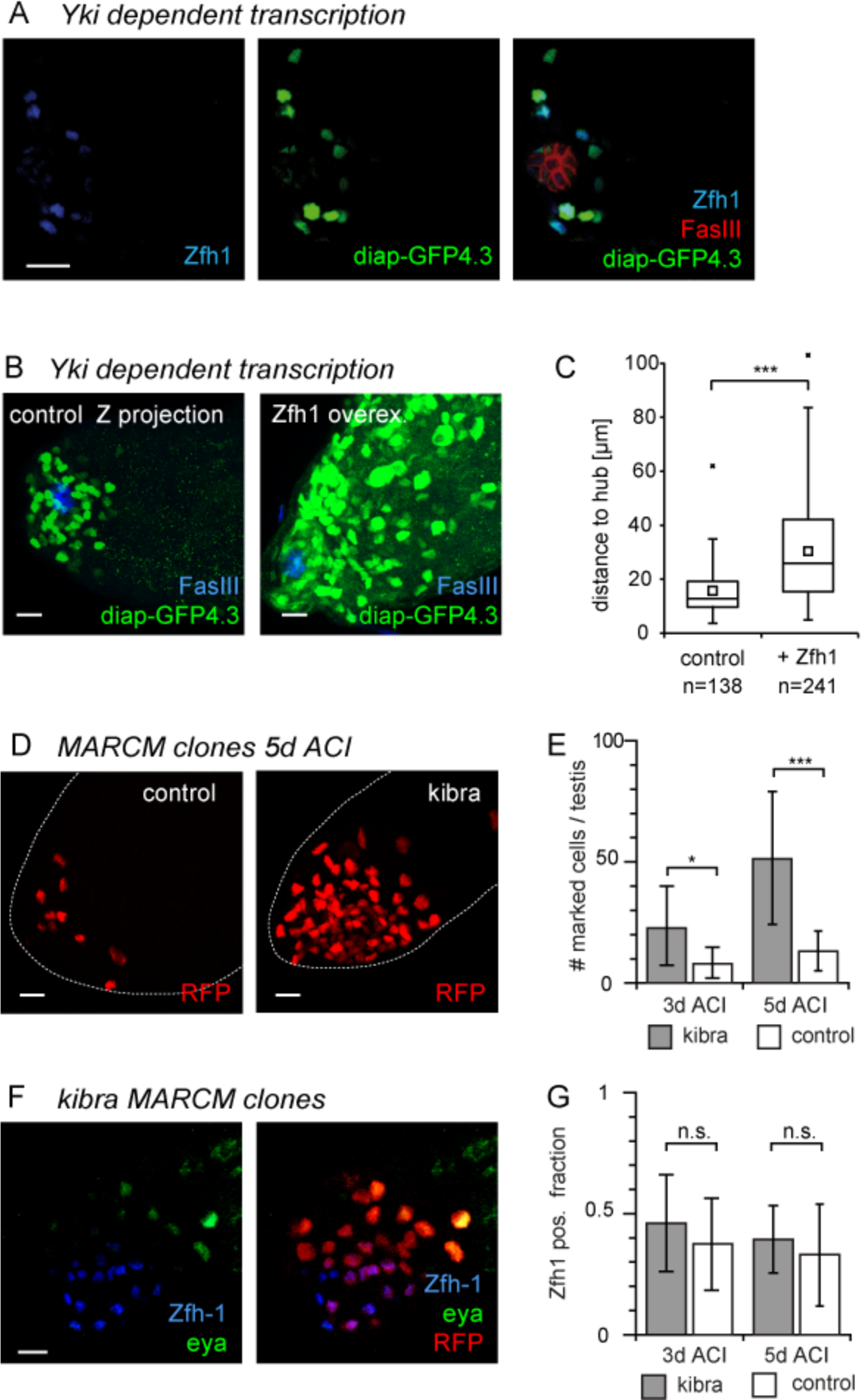
The Hippo pathway controls CySC proliferation. **A** Activity of a GFP transcriptional reporter (green) based on a Yki dependent fragment of the diap1 promoter is largely restricted to Zfh1 positive CySCs (blue). Hub marked by FasIII (red). **B** Zfh1 overexpression in the somatic lineage (8d) expands the population of cells with active Yki to regions further from the hub (FasIII, blue). **C** Quantification of **B** **D** Homozygous *kibra* mutant MARCM clones (RFP, red) expand relative to control clones. **E** Quantification of **D** **F)** A *kibra* clone (RFP, red) containing both cells expressing the stem cell marker Zfh1 (blue) or the differentiation marker Eya (green). **G** *kibra* and control clones do not differ in the fraction of Zfh1 positive clonal cells. Box indicates first and third quartile; horizontal line, median; square, mean; whiskers, data range up to 1.5x interquartile distance. Outliers shown individually. Bar graphs, mean ± SD. *, p<0.05 ***, p<0.01 (t-test), n.s., not significant. Scale bars 10*μ*m.

For the remaining experiments addressing the link betwen Zfh1 expression, Hpo signalling, and CySC proliferation we focussed on two of the putative target Zfh1 genes, *sav* and *kibra*.

### Sav and Kibra limit CySC proliferation

We first confirmed that *kibra* and *sav* affected proliferation in the somatic lineage of the testis, as previously shown for *hpo* (Amoyel et al., 2014). Homozygous *kibra* clones rapidly expanded relative to controls (Figure 4D,E). BrdU labelling confirmed that the expansion of the *kibra* clones was caused by increased proliferation (Figure S5C,D). Homozygosity for *kibra* did not block differentiation, as indicated by the expression of both the CySC marker Zfh1 and the CyC differentiation marker Eya (Fabrizio et al., 2003) in different cells of the same clone (Figure 4F). Both three and five days after clone induction (ACI) the fraction of Zfh1 positive cells did not differ between control and *kibra* mutant clones (Figure 4G). The increased proliferation of the *kibra* mutant stem cells is thus uncoupled from their ability and propensity to differentiate.

The effect of clonal inactivation of *sav* was less pronounced in the short term but was, as for *kibra*, readily detectable by increased long term retention of the clones, reflecting a reduced rate of loss from the niche due to skewed neutral competition (Amoyel et al., 2014) (Figure S5E,F).

### Kibra is expressed in the somatic lineage and is downregulated by ectopic Zfh1 expression

However, these experiments do not answer whether the putative niche signal - Zfh1 - Hippo axis regulates CySC proliferation under physiological conditions. We therefore wanted to know whether Kibra was expressed in the somatic lineage, and whether its expression was reduced in the Zfh1-positive CySCs. However, in the adult testis, Kibra antibody staining (Figure 5A) primarily marks the spectrosomes and fusomes in the germline cells, even though the Hpo pathway does not regulate the proliferation of these cells (Amoyel et al., 2014; Sun et al., 2008). In the somatic lineage, Kibra protein could with confidence only be observed in the hub, while the diffuse staining at the interface of germline and somatic cells could not be assigned to either cell type due to their tight apposition in the testis tip.

**Figure 5.**
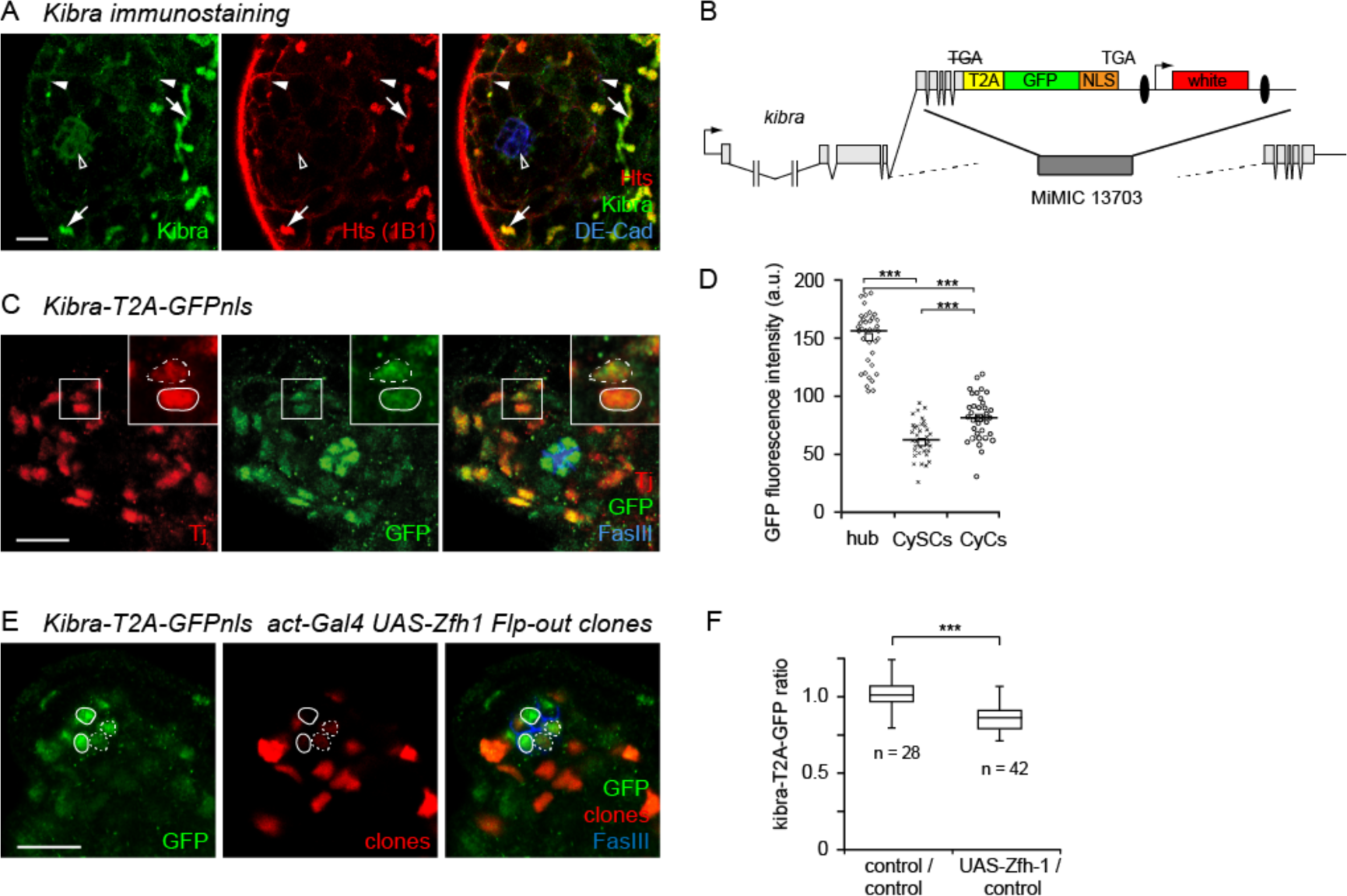
Kibra expression in the testis. **A** Kibra immunostaining (green) is detectable in the somatic hub cells (DE-Cad, blue, open arrowhead) and the spectrosomes and fusomes (Hts, red, arrows) within the germline. Diffuse, membrane associated staining around the cell outlines (solid arrowheads) cannot be assigned unambiguously to either lineage. **B** kibra-T2A-GFPnls construct inserted into a MiMIC landing site preceding exon 5. Ellipses, loxP sites. **C-F** GFP immunostaining in the Kibra-T2A-GFPnls reporter line. **C** Nuclear GFP is visible both in the germline (large, diffusely stained nuclei) and the somatic lineage, with pronounced staining in the hub (FasIII, blue). Note reduced signal in CySC nuclei identified as Tj-positive (red) nuclei abutting the hub (solid outline in inset) relative to their CyC neighbours (dashed outline). **D** Quantification of GFP immunostaining. **E** Zfh1 overexpression in RFP marked Flp-out clones (red, dashed outline) in the hub (FasIII, blue) decreases nucler GFP levels (green) relative to sibling cells (solid outline). **F** Ratio of GFP immunofluorescence between Zfh1 expressing and adjacent non-expressing nuclei or between controls. Scale bars 10*μ*m. Solid line, median; square, mean; ***, p<0.01 (ANOVA). Box indicates first and third quartile and median. Whiskers indicate data range up to 1.5x interquartile distance. ***, p<0.01 (t-test).

In contrast, the nuclei of the two lineages can be readily separated. To assess *kibra* transcription in the testis we therefore generated a construct containing *kibra* exons 5- 9, C-terminally fused to a nuclear GFP via a T2A site. This construct was then targeted to a MiMIC landing site (MI13703) (Venken et al., 2011) inserted in the fourth intron of the *kibra* locus (Figure 5B), resulting in viable and fertile kibra-T2AGFPnls flies. Nuclear GFP expression confirmed *kibra* transcription in both germline and soma. Within the somatic lineage nuclear GFP immunofluorescence was strongest in the hub (184% of median CyC levels) (Figure 5C,D). A somewhat weaker signal was also visible in CySCs, identified here as cells abutting the hub and expressing the somatic marker Tj, and in recently differentiated CyCs (Tj positive cells one tier further out) (Figure 5C). Compared with the CyCs, median nuclear GFP levels in the CySC were reduced by 23% (Figure 5D), consistent with the hypothesis that Zfh1 downregulates *kibra* transcription.

However, the presence of multiple Zfh1 DamID peaks near and within the *kibra* transcription unit (Figure S4) made it impractical to test directly whether *kibra* expression in the CySCs is downregulated by endogenous Zfh1 by deleting the corresponding binding sites. Conversely, clonal inactivation of Zfh1 causes the rapid exclusion of the mutant cells from the stem cell compartment (Leatherman and Dinardo, 2008), making it impossible to decide whether a possible effect on *kibra* expression would be directly caused by the absence of Zfh1, or be a secondary consequence of differentiation.

We therefore turned to mosaic expression of Zfh1 in the hub cells, that endogenously express Kibra but not Zfh1. This allowed us to test *in vivo* whether Zfh1 expression was sufficient to suppress *kibra* transcription. In line with the observations on the Zfh1 positive stem cells and their Zfh1 negative progeny, GFP levels in hub cell nuclei overexpressing Zfh1 were reduced by 15% relative to their non-overexpressing neighbours (Figure 5E,F).

### Sav expression is sensitive to endogenous Zfh1

Compared with *kibra*, the *sav* locus is smaller and possesses only a single Zfh1 DamID peak located 3′ of the coding region (Figure 3A). To test whether endogenous Zfh1 expression is sufficient for downregulating its target genes we concentrated on Sav, using a series of GFP based reporter constructs (Figure 6A). We isolated a genomic fragment spanning the *sav* genomic region, extending into the transcription units of the flanking genes and encompassing the entire Zfh1 DamID peak. As for Kibra, we tagged Sav with T2A-GFPnls. Nuclear GFP from this sav-T2A-GFPnls reporter was expressed in the germline, with a weaker signal in the somatic CySCs and CyCs, but, unlike *kibra*, no expression in the hub (Figure 6B). Derepression of *sav* in the CyCs could potentially negatively affect stem cell maintenance. Before modifying the presumptive control region we therefore deleted the Sav coding region and T2A cassette from our reporter (Figure 6A). The GFP expression pattern of this GFPnls-full_length construct was indistinguishable from that of the sav-T2A-GFPnls reporter (Figure 6B,C).

**Figure 6.**
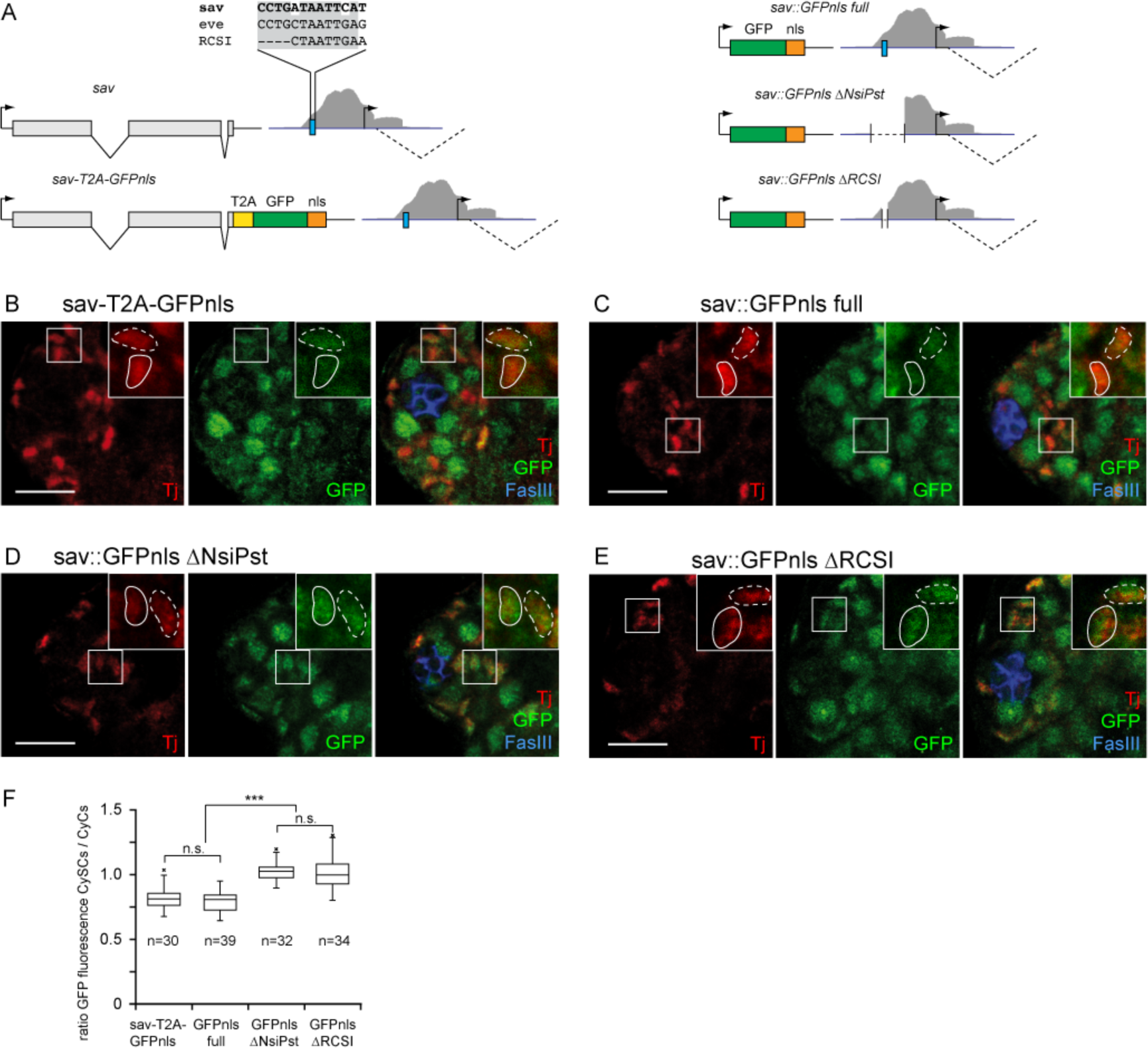
Sav in the adult testis. **A** Organization of the *sav* locus, the sav-T2A-GFPnls construct, and the sav::GFPnls transcriptional reporters. Blue line, extent of Zfh1 DamID peak (gray shading). Blue box, RCSI-like putative Zfh1 binding site. Sequences from *sav*, the *eve* Zfh1 binding site, and the original RCSI sequence are indicated. **B-E** GFP immunofluorescence from the *sav* reporter constructs. Note GFP signal (green) in the large germline nuclei and smaller somatic nuclei (marked by Tj, red); hub marked by FasIII (blue). **B** In the sav-T2A-GFPnls fusion construct, the GFP signal in CySC nuclei (identified by proximity to hub, solid outline) is reduced relative to the adjacent CyC nuclei (dashed outlines). **C** Same for the sav::GFPnls_full construct **D** In sav::GFPnls_ΔNsiP transgenic flies the difference in GFP intensity between CySC and CyC nuclei vanishes. **E** Same for the sav::GFPnls_ΔRCSI transgene. **F** Ratio of GFP signals in adjacent CySC and CyC nuclei for **B-E**. Scale bars 10*μ*m. Box indicates first and third quartile and median. Whiskers indicate data range up to 1.5x interquartile distance. Outliers marked individually; ***, p<0.01 (ANOVA); n.s., not significant.

DamID resolution is intrinsically limited by the distribution of the GATC target sites. In the case of the *sav* Zfh1 DamID peak, the flanking GATC sites are separated by 1063bp. This region contains a short motif with high similarity (10bp of 12bp identical) to the repressive Zfh1 homeodomain binding site within the *eve* mesodermal enhancer (Su et al., 1999), that had in turn been recognized due to its similarity to the conserved RCSI homeodomain binding motif used for cloning *zfh1* (Fortini et al., 1991). (Figure 6A). Similar, putative Zfh1 homeodomain binding sites corresponding to the degenerate sequence CTAATYRRNTT used to identify the *eve* RCSI-like motif (Su et al., 1999) were indeed significantly enriched in Zfh1 DamID peaks from both S2 cells and CySCs (S2 cells: PWMenrich raw score 1.377; p=8.06 E-11; CySCs: raw score 1.421; p=1.377 E-07; Stojnic, R. and Diez, D. (2015), PWMEnrich, R package 4.6.0).

We therefore first removed the central third of *sav*/*CG17119* intergenic region that overlaps with the Zfh1 DamID peak and includes the RCSI-like sequence motif (GFPnls-ΔNsiPst). In the final construct, we selectively deleted this putative Zfh1 binding site (GFPnls-ΔRCSI) (Figure 6A).

We then quantified relative GFP fluorescence in CySC and adjacent CyC nuclei for all four reporter lines (Figure 6B-F). For both the sav-T2A-GFPnls and the GFP-full length constructs (Figure 6B,C) GFP fluorescence was about 20% lower in the endogenously Zfh1 positive CySCs (again identified by position and Tj immunostaining) than in their differentiating CyC neighbours (Figure 6F). This difference was abolished for both constructs carrying the deletions within the Zfh1 DamID peak (GFPnls-ΔNsiPst and GFPnls-ΔRCSI) (Figure 6D-F), confirming that the RCSI-like motif is the *cis*-acting element responsible for downregulation of *sav* expression in the Zfh1-positive CySCs.

Taken together our observations show that the DamID experiments successfully uncovered target genes regulated by Zfh1 in the adult testis, that the regions flagged as Zfh1 binding peaks contain sequence elements conveying Zfh1 sensitivity, and that Zfh1 promotes CySC proliferation by downregulating transcription of Hippo pathway components.

## Discussion

Stemness and differentiation have traditionally largely been treated as alternative outcomes of a binary decision made by a stem cell that would globally govern its subsequent behaviour. Thus, the role of the niche would be to act upstream and bias this decision (Fuller and Spradling, 2007) (Figure 7A).

**Figure 7.**
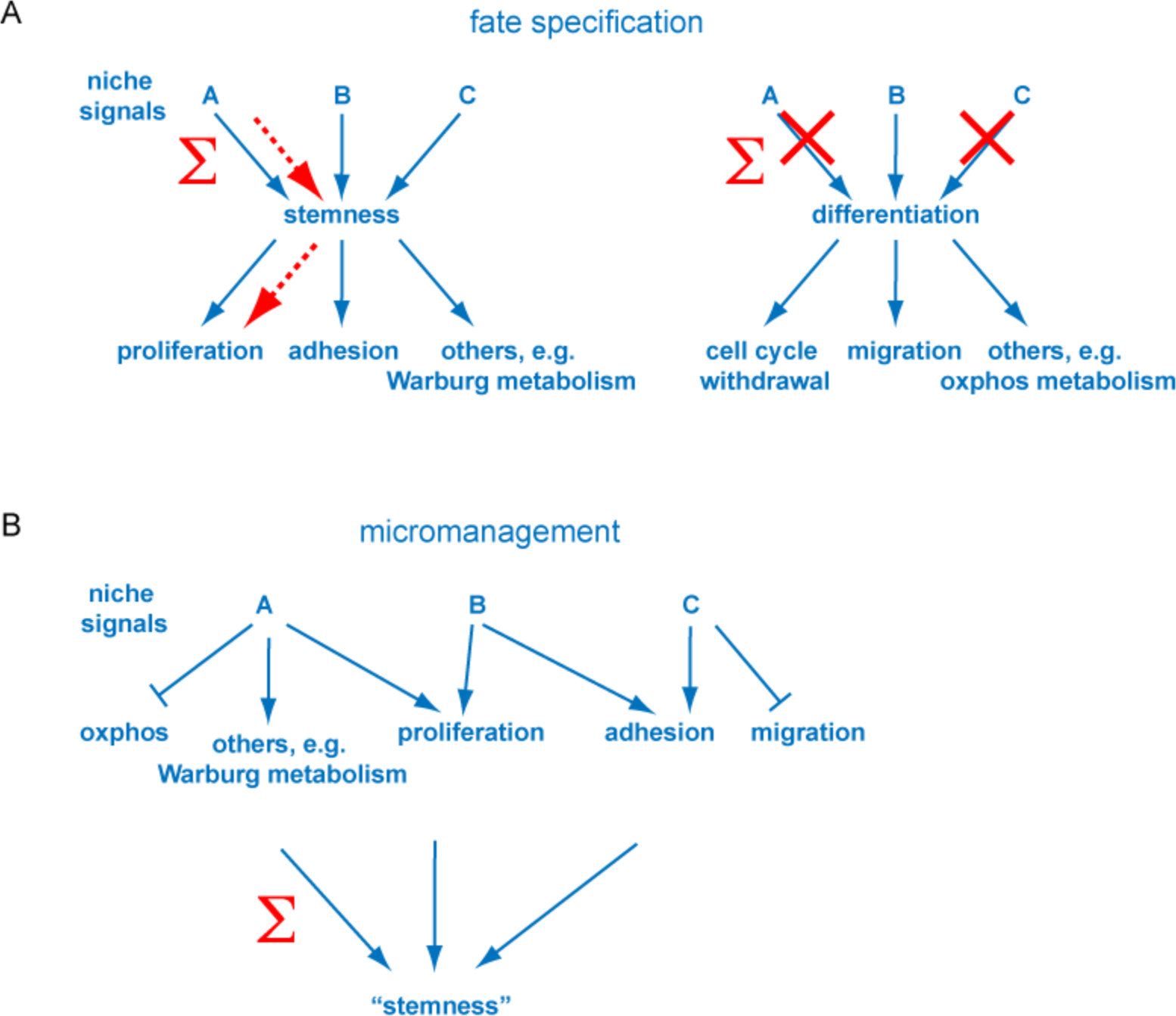
Models of stemness and niche function. **A** According to the traditional model, the various niche signals are integrated into a decision to adopt stem cell fate. Differentiation is the default response when such signals are absent. Explaining why increasing one particular niche signal impinges on only one or few aspects of stem cell behaviour (red, dashed arrows) is not straightforward. **B** Under the proposed micromanagement model, niche signals continuously control genetically separable subsets of stem cell behavioural output. Stemness becomes a compound phenotype rather than a clear cell fate.

However, we and others had previously shown that experimental activation of the Hh pathway upregulates proliferation of somatic CySCs, but does not affect the differentiation propensity of the mutant clones (Amoyel et al., 2013; Michel et al., 2012). Overexpression of Upd instead drives the proliferation of stem cell like cells that fail to differentiate but also partially acquire hub functions (Leatherman and Dinardo, 2008). Manipulation of MAPK signalling can uncouple stem cell self-renewal from the regulation of cell competition and adhesion to the niche (Amoyel et al., 2016; Singh et al., 2016). Manipulation of either of these endogenously active signalling pathways thus impinges on overlapping but distinct subsets of stem cell properties. However, such partial alteration of stem cell behaviour should not be possible if niche signals became integrated into a decision between stemness and differentiation.

The results presented here instead fill some crucial, mechanistic gaps to demonstrate a direct and unbroken regulatory link between niche signal input and stem cell proliferation. In the fly testis the Hh and Upd niche signals converge on Zfh1 (Amoyel et al., 2013; Leatherman and Dinardo, 2008; Michel et al., 2012), which we found here to bind to multiple genes encoding regulators of the Hippo signalling cascade. For two of these genes, *sav* and *kibra*, we could show that this results in a partial suppression of transcription, as reflected by reduced reporter gene expression in endogenously Zfh1 positive CySCs. For *sav* we were able to map the *cis*-acting sequence conveying this repression to a conserved, putative homeodomain binding motif within the Zfh1 DamID peak, while misexpression in the hub demonstrated that Zfh1 is sufficient to suppress *kibra* transcription *in vivo*. Even though both genes are downregulated in CySCs by no more than 25%, Yki activation in the testis is tightly restricted to the Zfh1 positive stem cells, with the small fraction of single positive cells presumably explained by differences in the maturation and degradation rates of the two proteins. Since the loss of one copy of any individual Hpo pathway gene has no obvious phenotype, we must postulate that Yki activity is controlled cooperatively through the modulation of multiple regulators that may, in turn, be influenced by multiple upstream stimuli: While *kibra* and *sav* appear to be controlled by niche signalling, the upstream Hippo pathway regulator Merlin has recently been implicated in regulating stem cell proliferation in response to cell contacts (Inaba et al., 2017). Surprisingly, in the ovary Hh regulates Hippo signalling through transcriptional control of Yki (Huang and Kalderon, 2014). At the moment we have no explanation as to why the male and female niches should make use of the same molecular players but follow a different regulatory logic. Nevertheless, our results show how in the testis one specific, genetically separable aspect of stem cell behaviour, i.e. proliferation, can be seamlessly traced back to niche signal input.

Generalizing from this direct control of proliferation, and considering other examples where different aspects of stem cell behaviour can be uncoupled at the level of niche signal input (Amoyel et al., 2016; Leatherman and Dinardo, 2008; Singh et al., 2016), we would like to suggest that the function of the niche is to directly instruct competent cells to execute specific stem cell behaviours. “Stemness” would thus become an operational description given to any cell successfully instructed to self renew and produce differentiating offspring, without having to invoke a cell fate decision. We would like to propose the term “micromanagement” for this mode of niche mediated stem cell regulation (Figure 7B).

Of course, this model comes with several caveats. First, the somatic CySCs of the fly testis or the stem cells of the mammalian intestinal epithelium may represent one extreme within a continuum of stem cell / niche systems that differ in the relative contributions of fate specification and micromanagement. The mammalian hematopoietic system, which has guided much of our thought about stemness and niches (Scadden, 2014), and indeed the *Drosophila* germline stem cells, which are maintained primarily through suppression of the differentiation factor Bam by a hub derived BMP niche signal (Chen and McKearin, 2003; Kawase et al., 2004) may represent the other end of the spectrum.

Second, the competence of a cell to respond to a given niche microenvironment does, to some extent, depend on cell fate decisions along its developmental trajectory: Unlike the CySC/CyC lineage, the muscle sheath of the testis does not respond to Upd or Hh overexpression by proliferating, even though both cell types are of mesodermal origin. However, in this context, the micromanagement model may provide a simple explanation for the phenomenon of lineage restriction observed in vertebrate regeneration models such as the zebrafish, a phenomenon that is again not readily compatible with treating stemness as the diametral opposite of differentiation (Sehring et al., 2016): If the transient niche formed after amputation did not induce actual dedifferentiation (i.e. a true cell fate decision establishing stem cell state), but cells in the stump were merely competent to respond to niche signals by temporarily suspending their normal physiological activities (e.g. ECM deposition in the case of osteoblasts) in favour of proliferating and migrating to the blastema, the need to postulate a separate cell fate memory disappears.

Third, the micromanagement model we propose will still be a truncation of the true regulatory network. For example, we already know that different aspects of stem cell activity in the testis are coupled by competition (Amoyel et al., 2016; Issigonis et al., 2009; Michel et al., 2012; Singh et al., 2016), making it difficult to analyze loss of function conditions, as affected cells are eliminated by differentiation regardless of the actual function of the affected pathway. In addition, the subsets of stem cell behaviours controlled by different niche signals may overlap, and any individual signal can control multiple aspects of stemness. For example, Upd not only cooperates with Hh to promote CySC proliferation through Zfh1, but independently ensures male specific development of the somatic lineage through Chinmo (Flaherty et al., 2010; Ma et al., 2014).

Even Zfh1 itself is likely to control additional aspects of stemness: Hippo pathway components account for only a small fraction of the many hundred potential target genes we identified in the CySCs. We found Zfh1 peaks e.g. also near multiple genes involved in glucose and energy metabolism, with carbohydrate transport appearing as a significantly enriched GO term. This invites the intriguing speculation that a potential, Warburg-like stem cell metabolic state (Chandel et al., 2016) may also be the outcome of detailed, continuous regulation by the niche rather than an intrinsic property of the stem cells.

Finally, under our micromanagement model, the minimal unit of stemness is a stem cell together with its niche. Obviously this can only hold true for tissue resident stem cells that actually are integrated with a niche, but may not apply to embryonic stem cells that are, at least *in vitro*, able to retain their stem cell properties in the absence of a complex signalling microenvironment.

## Materials and methods

### Fly stocks and transgenic lines

w;UAS-zfh1; w;;UAS-RedStinger; w;;FRT82B w^+^90E; w;Sco/CyO;tub-GAL80^ts^; y w;MiMIC kibra^MI13703^; y w;;Act5C>CD2Y>GAL4; and y sc v nos-phiC31Int;;attP2 were obtained from the Bloomington Drosophila Stock Center.

The following stocks were graciously provided by our colleagues: UAS-Hop^tumL^ (Bruce Edgar); y w vasa-Cas9/FM7c (Anne Morbach); UAS-LT3-Dam (Andrea Brand); ti Gal4 Gal80^ts^ (Doug Allan); / CyO, y^+^;FRT82 kibra^1^ / TM6B, Tb, Hu; y w;Sp/CyO,y^+^;diap1-GFP4.3/TM6B, Tb; w;FRTD 42D yki^B5^/CyO (Hugo Stocker); FRT 82B sav^3^/TM3 Sb (Florence Janody, Nic Tapon).

MARCM clones (Lee and Luo, 1999) were generated by crossing FRT males to w hs-FLP C587-Gal4 UAS-RedStinger virgins carrying the appropriate tub-Gal80 FRT chromosomes and heat shocking adult males for 1h at 37°C.

The zfh1-GFP and zfh1-T2A-Gal4 transgenes were generated by coinjecting two pU6-BbsI-gRNA plasmids (Gratz et al., 2014) (Addgene 45946) with a pRK2 (Huang et al., 2008) derived plasmid containing the transgene sequence, 1kb homology arms, and a w^+^ selection marker into y w vasa-Cas9/FM7c embryos.

The kibra-T2A-GFP line was generated by injecting a pBSKS-attB1_2SASD-0 plasmid (gift from Frank Schnorrer) containing kibra exons 5 to 9 fused to T2AGFPnls into nos-phiC31Int;MiMIC kibra^MI13703^ embryos.

The Sav reporter constructs were generated by fusing a T2A-GFPnls cassette to the Sav C-terminus in a genomic construct extending into the coding regions of the adjacent genes. Next, the sav coding region and the 3′ UTR segments were excised by fusion PCR and deletions introduced to the Zfh1 DamID peak. Constructs were cloned into pUAST-attB, removing the UAS sites, and injected into nos-phiC31Int;;attP40 embryos.

For UAS-LT3-zfh1-Dam the zfh1-RB ORF (Korneel Hens) was inserted into pUAST-attB-LT3-Dam (Andrea Brand) at Dam start codon and the plasmid injected into nos-phiC31Int;;attP2 embryos.

Full sequences are available upon request.

### Antibodies and immunohistochemistry

Testes were stained as described (Michel et al., 2011), incorporating an additional 1h permeabilization step with 1% Triton X-100 following fixation. The following antisera were used: rat-anti-DECadherin (DSHB, DCAD2) 1:100, mouse-anti-Eya (DSHB, eya10H6) 1:20, mouse-anti-FasIII (DSHB, 7G10) 1:100, rabbit-anti-GFP (Clontech) 1:500, rabbit-anti-Kibra (Nic Tapon) 1:200, rat-anti-Tj (Dorothea Godt) 1:250, Rabbit anti-vasa (Paul Lasko) 1:5000, rabbit-anti-Zfh1 (Ruth Lehmann) 1:4000. Secondary ABs raised in goat and labelled with Alexa-488, -568, or -633 (Invitrogen) were used 1:500.

### BrdU labeling

Flies were fed on food containing 2 mg/ml BrdU for 8h. In addition to the standard immunostaining protocol testes were incubated for 30min in 2M HCl, neutralized with 100mM borax solution and incubated with mouse anti-BrdU-Alexa 488 (BD) 1:200 over night.

### Imaging and image analysis

Images were acquired using Leica SP5, Zeiss LSM700, and Zeiss LSM780 confocal microscopes with 40x or 63x water immersion objectives. Unless stated otherwise, images are single optical slices. Image quantifications were performed using Fiji (Schindelin et al., 2012), applying a double blind protocol where appropriate. Differences between samples were tested for significance using Student’s t-test or ANOVA followed by Tukey’s HSD as appropriate. Images were prepared for publication using Adobe Photoshop and Illustrator.

### Cell culture

S2 cells were cultivated at room temperature in Schneider’s Drosophila Medium (Pan Biotech) with 10% FBS (Thermo Scientific) and transfected using a standard calcium phosphate protocol. Stable cell lines were selected on hygromycin B (Sigma).

### RNAseq

Zfh1 and Zfh1-CIDm expression was induced with 1mM CuSO4 for 24h before RNA extraction using Direct-zol (Zymo Research). RNA was quality checked with bioanalyser (Aligned Genomic) and sequenced by the CRTD/Biotec NGS facility using an Illumina HiSeq2000 machine.

### DamID

For the in vitro experiments DNA was extracted from stable mtn-Dam or mtn-Zfh1-Dam cell lines with Trizol (Sigmaaldrich) without Cu^2+^ induction. For the in vivo experiments DNA was extracted from testes proper of 50 zfh1-T2A-Gal4 tubGal80^ts^/UAS-LT3-Zfh1-Dam tub-Gal80^ts^ and zfh1-T2A-Gal4 tub-Gal80^ts^/UAS-LT3-Dam tub-Gal80^ts^ males following 24h induction at 30°C as described in (Laktionov et al., 2014). Samples were prepared according to (Southall et al., 2013) and sequenced by the CRTD/Biotec NGS facility using a Illumina HiSeq2000 machine.

### Bioinformatics

NGS reads from the DamID experiments were mapped to the Drosophila genome (dm6) using bowtie2. Zfh1 peaks were called using damid_seq analysis pipeline (Marshall and Brand, 2015). Results were analyzed using the find_peaks script (Marshall and Brand, 2015) with the false discovery rate (FDR) threshold set to 0.01. Overlap of the Zfh1 DamID peaks thus identified in S2 cells and CySCs with each other, Zfh1 ChIP peaks from Kc167 cells (Negre et al., 2011), or STARR-seq peaks from S2 cells or OSCs (Arnold et al., 2013) was analyzed using the peakPermTest resampling / permutation approach implemented in the ChIPpeakAnno bioconductor package (Zhu et al., 2010) excluding the ENCODE *Drosophila* blacklist (Celniker et al., 2009). For each comparison, we performed 1000 resampling runs artificially redistributing peaks while conserving their distribution of relative positions to landmarks of the respective, associated genes (transcription starts and end, introns, and coding regions). Overlap between associated genes (defined as genes < 1kb distant from the ends of a Zfh1 DamID peak or other feature) was tested for significance using the *χ*^2^-test.

Enrichment of the degenerate, RCSI-like motif (CTAATYRRNTT) within Zfh1 DamID peaks was detected with help of the PWMenrich package (Stojnic, R. and Diez, D. (2015). PWMEnrich, R package).

RNASeq analysis was performed using the DESeq R package (Anders and Huber, 2010). Reads were normalized according to the library complexity. Gene annotation was obtained from ENSEMBL (BDGP6 version) using the biomaRt R package (Smedley et al., 2009). Genes were flagged as significantly changing expression if the corresponding adjusted p-values were below 0.05.

## Acknowledgements

We would like to thank the colleagues mentioned individually above and the JEDI community for fly stocks, reagents, and advice, the CRTD NGS sequencing facility for their support, Anastasia Labudina and Ilker Deniz for their help with immunostainings, and Heiner Grandel, Christian Lange, Nikolay Ninov, and Pavel Tomancak for discussions and critical reading of the manuscript. The project was supported by a CRTD seed grant and Deutsche Forschungsgemeinschaft (DFG) grant BO 3270/4-1 to CB.

## Author contributions

OAP performed the fly experiments, EAA performed the cell culture, DamID, and RNAseq experiments, EAA and NVT performed the bioinformatic analysis. All authors analysed and interpreted the data. CB initiated the project and wrote the manuscript.

## Data availability

DamID and RNAseq data will be placed in the appropriate repositories upon acceptance of the manuscript.

## References

Amoyel, M., Anderson, J., Suisse, A., Glasner, J. and Bach, E. A. (2016). Socs36E Controls Niche Competition by Repressing MAPK Signaling in the Drosophila Testis. PLoS Genet 12, e1005815.

Amoyel, M., Sanny, J., Burel, M. and Bach, E. A. (2013). Hedgehog is required for CySC self-renewal but does not contribute to the GSC niche in the Drosophila testis. Development 140, 56-65.

Amoyel, M., Simons, B. D. and Bach, E. A. (2014). Neutral competition of stem cells is skewed by proliferative changes downstream of Hh and Hpo. EMBO J 33, 2295-313.

Anders, S. and Huber, W. (2010). Differential expression analysis for sequence count data. Genome Biol 11, R106.

Arnold, C. D., Gerlach, D., Stelzer, C., Boryn, L. M., Rath, M. and Stark, A. (2013). Genome-wide quantitative enhancer activity maps identified by STARR-seq. Science 339, 1074-7.

Baumgartner, R., Poernbacher, I., Buser, N., Hafen, E. and Stocker, H. (2010). The WW domain protein Kibra acts upstream of Hippo in Drosophila. Dev Cell 18, 309-16.

Broihier, H. T., Moore, L. A., Van Doren, M., Newman, S. and Lehmann, R. (1998). zfh-1 is required for germ cell migration and gonadal mesoderm development in Drosophila. Development 125, 655-66.

Celniker, S. E., Dillon, L. A., Gerstein, M. B., Gunsalus, K. C., Henikoff, S., Karpen, G. H., Kellis, M., Lai, E. C., Lieb, J. D., MacAlpine, D. M. et al. (2009). Unlocking the secrets of the genome. Nature 459, 927-30.

Chandel, N. S., Jasper, H., Ho, T. T. and Passegue, E. (2016). Metabolic regulation of stem cell function in tissue homeostasis and organismal ageing. Nat Cell Biol 18, 823-32.

Chen, D. and McKearin, D. (2003). Dpp signaling silences bam transcription directly to establish asymmetric divisions of germline stem cells. Curr Biol 13, 1786-91.

Enderle, L. and McNeill, H. (2013). Hippo gains weight: added insights and complexity to pathway control. Sci Signal 6, re7.

Fabrizio, J. J., Boyle, M. and DiNardo, S. (2003). A somatic role for eyes absent (eya) and sine oculis (so) in Drosophila spermatocyte development. Dev Biol 258, 117-28.

Flaherty, M. S., Salis, P., Evans, C. J., Ekas, L. A., Marouf, A., Zavadil, J., Banerjee, U. and Bach, E. A. (2010). chinmo is a functional effector of the JAK/STAT pathway that regulates eye development, tumor formation, and stem cell self-renewal in Drosophila. Dev Cell 18, 556-68.

Fortini, M. E., Lai, Z. C. and Rubin, G. M. (1991). The Drosophila zfh-1 and zfh-2 genes encode novel proteins containing both zinc-finger and homeodomain motifs. Mech Dev 34, 113-22.

Fuller, M. T. and Spradling, A. C. (2007). Male and female Drosophila germline stem cells: two versions of immortality. Science 316, 402-4.

Genevet, A., Wehr, M. C., Brain, R., Thompson, B. J. and Tapon, N. (2010). Kibra is a regulator of the Salvador/Warts/Hippo signaling network. Dev Cell 18, 300-8.

Gheldof, A., Hulpiau, P., van Roy, F., De Craene, B. and Berx, G. (2012). Evolutionary functional analysis and molecular regulation of the ZEB transcription factors. Cell Mol Life Sci 69, 2527-41.

Gratz, S. J., Ukken, F. P., Rubinstein, C. D., Thiede, G., Donohue, L. K., Cummings, A. M. and O’Connor-Giles, K. M. (2014). Highly specific and efficient CRISPR/Cas9-catalyzed homology-directed repair in Drosophila. Genetics 196, 961-71.

Huang, J. and Kalderon, D. (2014). Coupling of Hedgehog and Hippo pathways promotes stem cell maintenance by stimulating proliferation. J Cell Biol 205, 325-38.

Huang, J., Zhou, W., Watson, A. M., Jan, Y. N. and Hong, Y. (2008). Efficient ends-out gene targeting in Drosophila. Genetics 180, 703-7.

Inaba, M., Buszczak, M. and Yamashita, Y. M. (2015). Nanotubes mediate niche-stem-cell signalling in the Drosophila testis. Nature 523, 329-32.

Inaba, M., Sorenson, D. R., Kortus, M., Salzmann, V. and Yamashita, Y. M. (2017). Merlin is required for coordinating proliferation of two stem cell lineages in the Drosophila testis. Sci Rep 7, 2502.

Irvine, K. D. and Harvey, K. F. (2015). Control of organ growth by patterning and hippo signaling in Drosophila. Cold Spring Harb Perspect Biol 7.

Issigonis, M., Tulina, N., de Cuevas, M., Brawley, C., Sandler, L. and Matunis, E. (2009). JAK-STAT signal inhibition regulates competition in the Drosophila testis stem cell niche. Science 326, 153-6.

Kawase, E., Wong, M. D., Ding, B. C. and Xie, T. (2004). Gbb/Bmp signaling is essential for maintaining germline stem cells and for repressing bam transcription in the Drosophila testis. Development 131, 1365-75.

Kiger, A. A., Jones, D. L., Schulz, C., Rogers, M. B. and Fuller, M. T. (2001). Stem cell self-renewal specified by JAK-STAT activation in response to a support cell cue. Science 294, 2542-5.

Lai, Z. C., Fortini, M. E. and Rubin, G. M. (1991). The embryonic expression patterns of zfh-1 and zfh-2, two Drosophila genes encoding novel zinc-finger homeodomain proteins. Mech Dev 34, 123-34.

Laktionov, P. P., White-Cooper, H., Maksimov, D. A. and Beliakin, S. N. (2014). [Transcription factor comr acts as a direct activator in the genetic program controlling spermatogenesis in D. melanogaster]. Mol Biol (Mosk) 48, 153-65.

Leatherman, J. L. and Dinardo, S. (2008). Zfh-1 controls somatic stem cell self-renewal in the Drosophila testis and nonautonomously influences germline stem cell self-renewal. Cell Stem Cell 3, 44-54.

Leatherman, J. L. and Dinardo, S. (2010). Germline self-renewal requires cyst stem cells and stat regulates niche adhesion in Drosophila testes. Nat Cell Biol 12, 806-11.

Lee, T. and Luo, L. (1999). Mosaic analysis with a repressible cell marker for studies of gene function in neuronal morphogenesis. Neuron 22, 451-61.

Losick, V. P., Morris, L. X., Fox, D. T. and Spradling, A. (2011). Drosophila stem cell niches: a decade of discovery suggests a unified view of stem cell regulation. Dev Cell 21, 159-71.

Ma, Q., Wawersik, M. and Matunis, E. L. (2014). The Jak-STAT target Chinmo prevents sex transformation of adult stem cells in the Drosophila testis niche. Dev Cell 31, 474-86.

Marshall, O. J. and Brand, A. H. (2015). damidseq_pipeline: an automated pipeline for processing DamID sequencing datasets. Bioinformatics 31, 3371-3.

Meng, Z., Moroishi, T. and Guan, K. L. (2016). Mechanisms of Hippo pathway regulation. Genes Dev 30, 1-17.

Mi, H., Poudel, S., Muruganujan, A., Casagrande, J. T. and Thomas, P. D. (2016). PANTHER version 10: expanded protein families and functions, and analysis tools. Nucleic Acids Res 44, D336-42.

Michel, M., Kupinski, A. P., Raabe, I. and Bokel, C. (2012). Hh signalling is essential for somatic stem cell maintenance in the Drosophila testis niche. Development 139, 2663-9.

Michel, M., Raabe, I., Kupinski, A. P., Perez-Palencia, R. and Bökel, C. (2011). Local BMP receptor activation at adherens junctions in the Drosophila germline stem cell niche. Nat Commun 2, 415.

Negre, N., Brown, C. D., Ma, L., Bristow, C. A., Miller, S. W., Wagner, U., Kheradpour, P., Eaton, M. L., Loriaux, P., Sealfon, R. et al. (2011). A cis-regulatory map of the Drosophila genome. Nature 471, 527-31.

Port, F., Chen, H. M., Lee, T. and Bullock, S. L. (2014). Optimized CRISPR/Cas tools for efficient germline and somatic genome engineering in Drosophila. Proc Natl Acad Sci U S A 111, E2967-76.

Postigo, A. A. and Dean, D. C. (1999). ZEB represses transcription through interaction with the corepressor CtBP. Proc Natl Acad Sci U S A 96, 6683-8.

Scadden, D. T. (2014). Nice neighborhood: emerging concepts of the stem cell niche. Cell 157, 41-50.

Schindelin, J., Arganda-Carreras, I., Frise, E., Kaynig, V., Longair, M., Pietzsch, T., Preibisch, S., Rueden, C., Saalfeld, S., Schmid, B. et al. (2012). Fiji: an open-source platform for biological-image analysis. Nat Methods 9, 676-82.

Sehring, I. M., Jahn, C. and Weidinger, G. (2016). Zebrafish fin and heart: what’s special about regeneration? Curr Opin Genet Dev 40, 48-56.

Singh, S. R., Liu, Y., Zhao, J., Zeng, X. and Hou, S. X. (2016). The novel tumour suppressor Madm regulates stem cell competition in the Drosophila testis. Nat Commun 7, 10473.

Smedley, D., Haider, S., Ballester, B., Holland, R., London, D., Thorisson, G. and Kasprzyk, A. (2009). BioMart--biological queries made easy. BMC Genomics 10, 22.

Southall, T. D., Gold, K. S., Egger, B., Davidson, C. M., Caygill, E. E., Marshall, O. J. and Brand, A. H. (2013). Cell-type-specific profiling of gene expression and chromatin binding without cell isolation: assaying RNA Pol II occupancy in neural stem cells. Dev Cell 26, 101-12.

Su, M. T., Fujioka, M., Goto, T. and Bodmer, R. (1999). The Drosophila homeobox genes zfh-1 and even-skipped are required for cardiac-specific differentiation of a numb-dependent lineage decision. Development 126, 3241-51.

Sun, S., Zhao, S. and Wang, Z. (2008). Genes of Hippo signaling network act unconventionally in the control of germline proliferation in Drosophila. Dev Dyn 237, 270-5.

Szymczak, A. L., Workman, C. J., Wang, Y., Vignali, K. M., Dilioglou, S., Vanin, E. F. and Vignali, D. A. (2004). Correction of multi-gene deficiency in vivo using a single ‘self-cleaving’ 2A peptide-based retroviral vector. Nat Biotechnol 22, 589-94.

Tapon, N., Harvey, K. F., Bell, D. W., Wahrer, D. C., Schiripo, T. A., Haber, D. and Hariharan, I. K. (2002). salvador Promotes both cell cycle exit and apoptosis in Drosophila and is mutated in human cancer cell lines. Cell 110, 467-78.

Tulina, N. and Matunis, E. (2001). Control of stem cell self-renewal in Drosophila spermatogenesis by JAK-STAT signaling. Science 294, 2546-9.

Venken, K. J., Schulze, K. L., Haelterman, N. A., Pan, H., He, Y., Evans-Holm, M., Carlson, J. W., Levis, R. W., Spradling, A. C., Hoskins, R. A. et al. (2011). MiMIC: a highly versatile transposon insertion resource for engineering Drosophila melanogaster genes. Nat Methods 8, 737-43.

Xie, T. and Spradling, A. C. (2000). A niche maintaining germ line stem cells in the Drosophila ovary. Science 290, 328-30.

Yu, J., Zheng, Y., Dong, J., Klusza, S., Deng, W. M. and Pan, D. (2010). Kibra functions as a tumor suppressor protein that regulates Hippo signaling in conjunction with Merlin and Expanded. Dev Cell 18, 288-99.

Zhang, L., Ren, F., Zhang, Q., Chen, Y., Wang, B. and Jiang, J. (2008). The TEAD/TEF family of transcription factor Scalloped mediates Hippo signaling in organ size control. Dev Cell 14, 377-87.

Zhu, L. J., Gazin, C., Lawson, N. D., Pages, H., Lin, S. M., Lapointe, D. S. and Green, M. R. (2010). ChIPpeakAnno: a Bioconductor package to annotate ChIP-seq and ChIP-chip data. BMC Bioinformatics 11, 237.

